# Heterogeneous signaling pathways are critical for the persistence of memory T cells in spleen and bone marrow

**DOI:** 10.64898/2026.04.02.714263

**Authors:** Emilia Schneider Revueltas, Lukas Almes, Koji Tokoyoda, Xiangyi Deng, Anna Casanovas Subirana, Marta Ferreira-Gomes, Rebecca Cornelis, Jun Dong, Frederik Heinrich, Pawel Durek, Mir-Farzin Mashreghi, Hyun-Dong Chang, Andreas Radbruch

## Abstract

Persistence of memory T lymphocytes, in the apparent absence of antigen, is a hallmark of immune memory and key to adaptive immunity to recurrent infections. The signaling pathways ensuring survival and quiescence of the memory T cells are largely enigmatic. Here we show, by inhibition *in vivo*, that persistence of surface CD69+KLF2-tissue-resident memory T cells of murine bone marrow and spleen is blocked by antibodies to the integrins VLA-4 and LFA-1, connecting the memory T cells to VCAM1 and ICAM1 of stromal cells. Persistence requires the PI3K/AKT signaling pathway, since it is blocked by Wortmannin, and it involves PI3K-dependent survival genes. Surface CD69-KLF2+ memory T cells of the bone marrow are also dependent on integrin-mediated contact to stromal cells. Their persistence critically depends on the NF-kB pathway, their PI3K signaling pathway is not relevant. Blocking Jak1 and 3 of the interleukin-7 and -15 signaling pathways does affect memory T cells of the spleen, but not those of the bone marrow. Thus, tissue-resident KLF2+ and KLF2-memory T cells, and memory T cells of spleen and bone marrow, use different signaling pathways, adapting them to their respective tissues and reflecting an unexpected heterogeneity in the molecular mechanisms of persistence.

## Introduction

The unique capacity of the adaptive immune system to memorize antigens once experienced is based on the long-term maintenance of antigen-specific “memory” T and B lymphocytes and long-lived plasma cells secreting antigen-specific antibodies (1,2) in the apparent absence of the antigen. For long-lived “memory” plasma cells, it has been shown that they are maintained in the bone marrow by integrin-mediated contact to stromal cells and BAFF or APRIL signal (3–5). The contact to stromal cells activates the Phosphatidyl-inositol-3-kinases (PI3K), Protein kinase B (Akt), inactivates the Forkhead box class O1 and 3a (FoxO1/3a) transcription factors and prevents activation of caspase 3, i.e. provides resilience to metabolic stress. BAFF or APRIL activate the nuclear factor kappa light chain enhancer of activated B cells (NF-kB) signaling pathway, prevent activation of caspase 12, i.e. resilience to anabolic stress(6).

The bone marrow does not only contain memory plasma cells, but also significant populations of memory B (7) and CD4 and CD8 memory T lymphocytes of systemic immunity (8,9). These memory T cells are maintained individually in contact to stromal cells expressing interleukin-7, and they rest in terms of proliferation, unless they are reactivated by antigen (10). A significant proportion of them expresses CD69 on the cell surface, a putative hallmark of tissue-resident memory T cells and their transcriptomic signature (11). Recently we have shown that also memory T cells of the bone marrow not expressing CD69 on the cell surface are residents of the bone marrow. The transcription factor Krüppel-like factor 2 (KLF2) is expressed by the surface CD69-negative, but not by the surface CD69-positive memory T cells (8,9). KLF2 itself is known as a transcription factor inducing quiescence and impacting on survival of T lymphocytes (12,13).

The uniform distribution of all CD4 and CD8 memory T cells in the bone marrow, their contact to stromal cells, and their proliferative rest (14–16) all alike memory plasma cells, provoke the question, whether these memory cells may use the same signaling pathways involved in the maintenance of memory plasma cells, i.e. the PI3K/Akt and the NF-kB pathways. And question whether the Jak/Stat signaling pathway, induced by interleukin-7 and -15, and required for “homeostatic proliferation”, is involved at all.

Here, we have tested the possible involvement and critical relevance of the PI3K/Akt, NF-kB and Jak1/3 signaling pathways, and the relevance of integrin-mediated binding to stromal cells, for the persistence of endogenous memory CD4 and CD8 lymphocytes of murine spleen and bone marrow, by blocking them *in vivo*, with integrin-specific antibodies, Wortmannin, IKK16 or Tofacitinib. Persistence of memory T cells in the bone marrow is blocked by VLA-4 and LFA1-specific antibodies. While surface CD69+ memory T cell persistence critically depends on PI3K signaling, surface CD69-memory T cells rather depend on NF-kB signaling in the bone marrow and their PI3K pathway is negligible for their persistence. Bone marrow memory T cells are not affected by inhibition of Jak1/3 signaling, while some of them in the spleen are. These results reveal an intriguing heterogeneity of mechanisms maintaining T cell memory, beyond homeostatic, cytokine-dependent proliferation.

## Results

To induce CD4 and CD8 memory T cells residing in spleen and bone marrow, C57BL/6 mice were immunized i.p. with 4-hydroxy-3-nitrophenylacetyl coupled chicken gamma globulin (NP-CGG) in incomplete Freund’s adjuvant on days 0, 21 and 42. 32 days later, on days 74, 76 and 78, the mice were treated with prospective inhibitors of molecular maintenance pathways, and sacrificed on day 80 (Fig. 1A). The numbers of surface CD69+ and surface CD69-, CD4+ and CD8+ memory T cells in the spleen and in the bone marrow were determined by flow cytometry (suppl. Fig. 1A). The inhibitors Wortmannin, IKK16 and Tofacitinib were titrated in the *in vivo* setting upfront (suppl. Figure 3).

**Figure 1:**
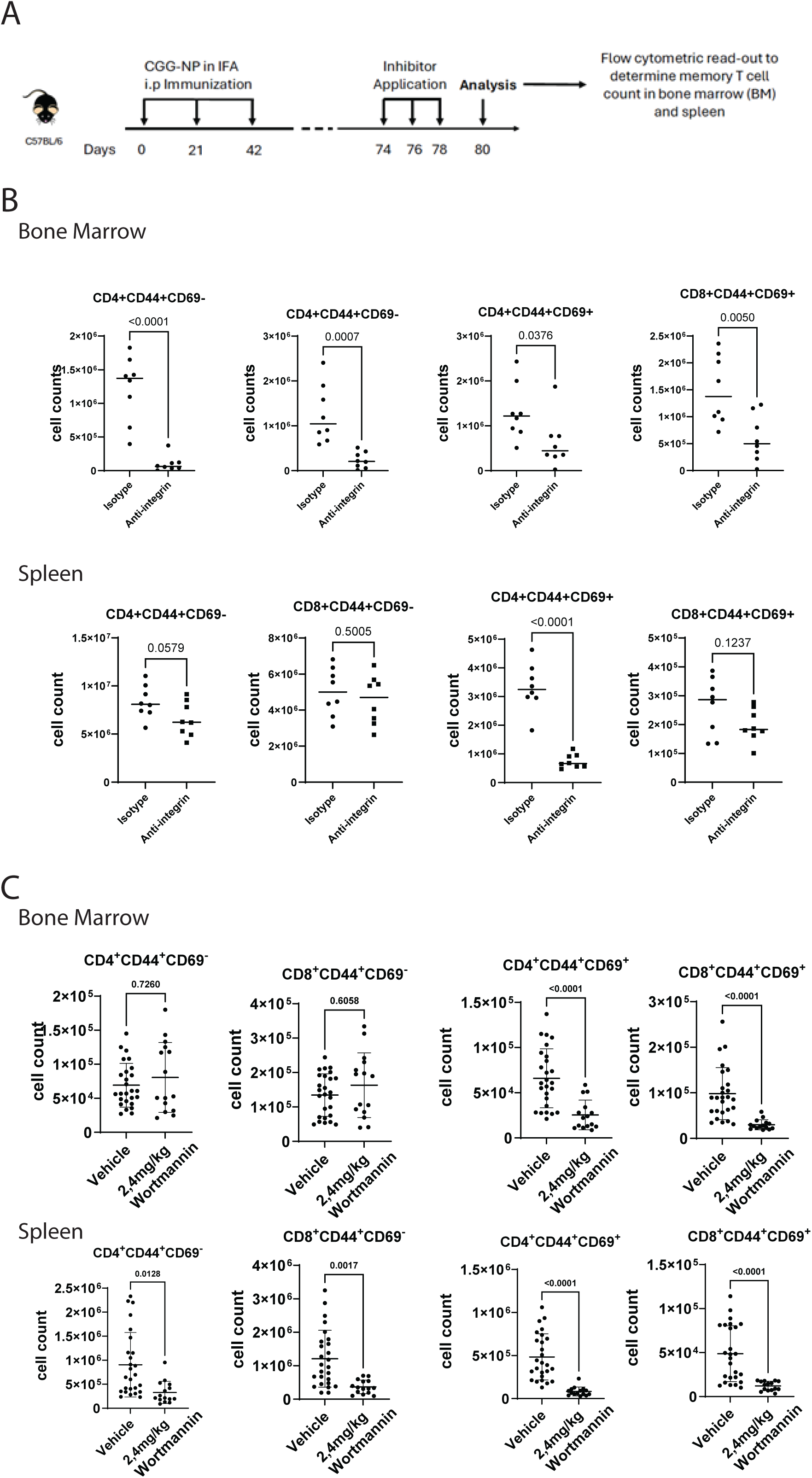
Experimental setup of *in vivo* inhibition of signaling pathways (A). Treatment of mice with antibodies to integrins VLA-4 and LFA-1, 200 μg/mouse each, and cytometric cell counts of mice treated with anti-integrins or isotype control antibodies (B). Treatment of mice with 2.4 mg/kg body weight of Wortmannin or vehicle (C). Each dot represents one mouse. Statistical analysis was performed in Prism 10 with an unpaired T-test.

### Integrin-mediated persistence of memory T cells

As previously used for the ablation of long-lived plasma cells from the bone marrow (5) we here used 200 μg anti-CD49d (integrin α4) and 200 μg anti-CD11a (integrin αL), or 200 μg rat IgG2a and 200 μg rat IgG2b isotype control antibodies per injection and animal, to block the VLA-4 (integrin α4β1) and LFA-1 (integrin αLβ2) links of memory T cells to VCAM1 (CD106) and ICAM1 (CD54) of the stromal cells they adhere to in the bone marrow (5). In the bone marrow of the treated mice, the simultaneous block of VLA-4 and LFA-1 resulted in significant ablation of CD4 and CD8, surface CD69+ and surface CD69-memory T cells, while in the spleen, only surface CD69+ CD4 memory T cells were significantly ablated (Fig. 1B). As expected, also plasma cells were depleted from bone marrow and spleen (suppl. Fig. 1B), while total cell numbers were not affected (suppl. Fig. 1C).

For long-lived (memory) plasma cells of the bone marrow, we had shown before that contact to stromal cells activates the PI3K pathway which promotes resilience to metabolic stress (6) Here we used Wortmannin to block PI3K activation in resting memory T cells *in vivo*. Interestingly, although both surface CD69+ and CD69-memory T cells are ablated from bone marrow by blocking the integrins VLA-4 and LFA-1, only the persistence of surface CD69+ memory T cells was blocked by Wortmannin. While in the spleen, persistence of all memory T cells subpopulations was blocked significantly (Fig. 1C) by inhibition of PI3K. It should be noted that absolute cell numbers and those of CD106+ (VCAM1) stromal cells were not affected by Wortmannin in the bone marrow, at the concentrations used (suppl. Fig. 1C).

In summary, these data suggest that integrin-induced PI3K signaling is critical for the persistence of surface CD69+, CD4 and CD8 memory T cells in the bone marrow, and for memory T cells in the spleen. Interestingly, persistence of surface CD69-negative memory T cells of the bone marrow is integrin-dependent but does not depend on PI3K signaling.

### Memory T cell persistence involving NF-kB and Jak/Stat signaling

To interrogate alternative signaling pathways possibly involved in the persistence of memory T cells, we also blocked NF-kB- and JAK/STAT signaling. We used IKK-16 to block NF-kB signaling. At a dose of 20 mg per kg body weight, IKK-16 significantly blocked the persistence of surface CD69-memory T cells in the bone marrow, but not in the spleen. The effects on persistence of surface CD69+ memory T cells were not significant (Fig. 2A). To block signaling of Janus kinases (JAK), including JAK3, which is the tyrosine kinase signaling downstream of the receptors for interleukin-7 and interleukin-15, we used Tofacitinib. At 20 mg per kg body weight, persistence of memory T cells of the spleen was significantly affected, although not completely blocked. Numbers of memory T cells of the bone marrow were not significantly affected, neither those of surface CD69+ nor those of surface CD69-memory cells (Fig. 2B).

**Figure 2:**
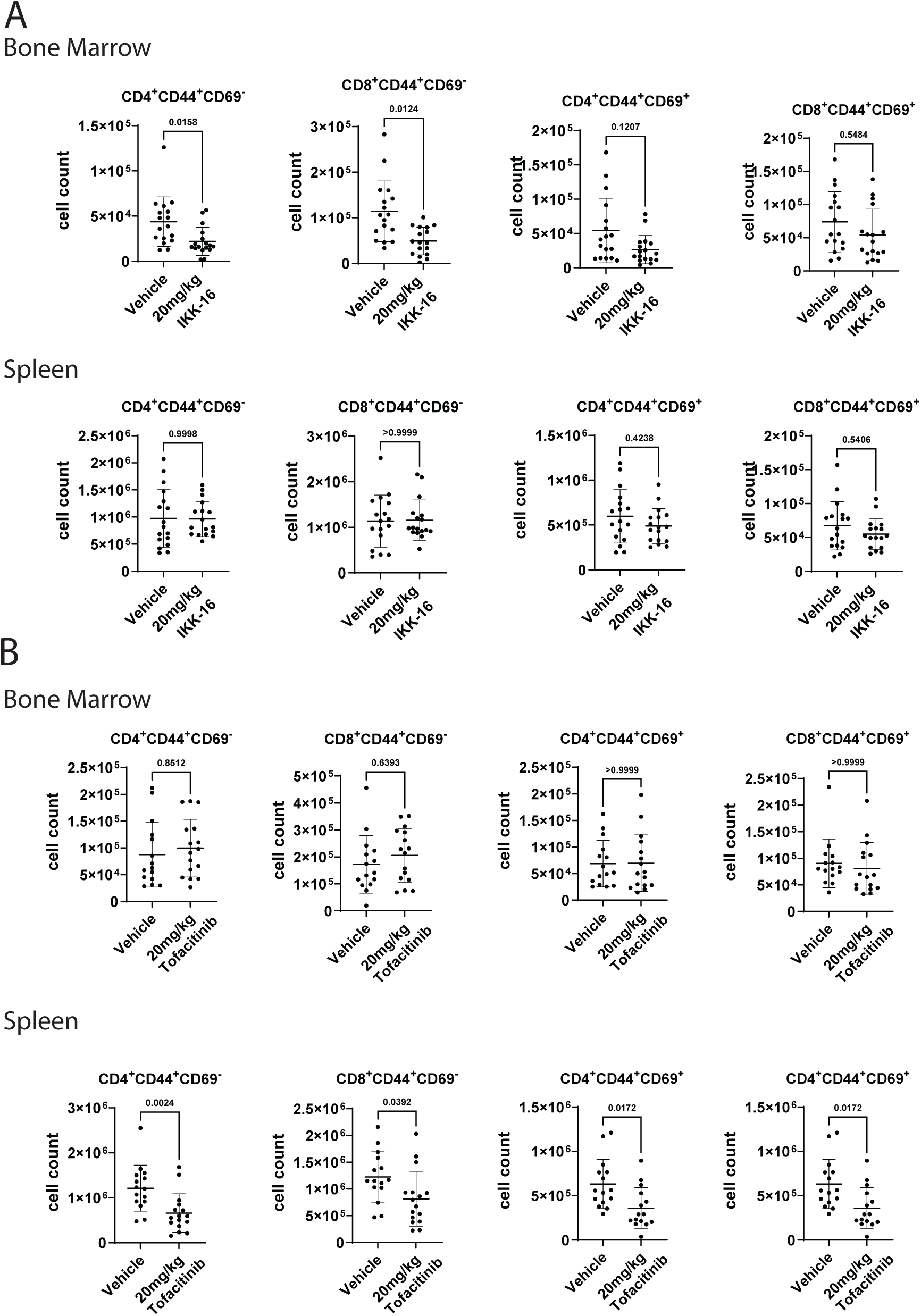
Treatment of mice with 20 mg/kg body weight of IKK16 (A), or with 20 mg/kg body weight of Tofacitinib (B) to block the NF-kB and JAK/STAT signaling pathways. Statistical analysis was performed in Prism 10 with an unpaired T-test.

Taken together, the data indicate that persistence of surface CD69+ memory T cells in bone marrow and spleen does not critically depend on NF-kB signaling. However, persistence of surface CD69-memory T cells of the bone marrow, but not of the spleen does. Blocking JAK/STAT signaling does affect memory T cell numbers in the spleen, but not in the bone marrow.

### PI3K is activated in resting KLF2-negative, surface CD69+ memory T cells of bone marrow

To determine PI3K pathway activity in Wortmannin-sensitive surface CD69+ and Wortmannin-refractory surface CD69-memory T cells, we sequenced the single cell transcriptomes based on surface CD69 CITE-seq expression of memory T cells of bone marrow of mice treated with Wortmannin or not. We previously reported for human memory T cells, that both surface CD69-negative and surface CD69-positive bone marrow memory T cells do transcribe and translate *CD69* to a similar extent. However, CD69-positive cells expressing it on the cell surface do not express the receptor 1 for sphingosin-1-phosphate, nor the transcription factor inducing it, which is KLF2 (8,17). In the present transcriptome analysis of resting CD8 memory T cells from bone marrow, cells were classified into 6 clusters using harmony-based integration of the different mice treated and untreated with Wortmannin (Fig. 3A-C). Cells of cluster 2 were most sensitive to Wortmannin (Fig. 3B). According to their transcriptomes, we classified them as memory T cells expressing a series of “innate” receptors, as is evident from the list of significant signature genes of the cluster (Fig. 3C), expression of which they share with NK cells. Some of them express KLF2, most do not (Fig. 3D). We then divided CD8 memory T cells of cluster 2 into KLF2-negative, surface CD69-positive and KLF2-positive, surface CD69-negative populations, and compared the transcriptomes of these cells in response to Wortmannin treatment (Fig. 3D-E). KLF2-positive, surface CD69-negative cells of cluster 2 do not show any difference in their global gene expression, when treated with Wortmannin, indicating that their PI3K signaling pathway is not active (Fig. 3F). For KLF2-negative, surface CD69-positive cells of cluster 2, the transcriptome is significantly affected by Wortmannin inhibition. This indicates that PI3K signaling is active, while they are resting and persisting, and revealing sets of genes up- and downregulated by PI3K (suppl. Figure 2). The precise role of these genes in promoting persistence of the memory T cells in the bone marrow remains to be clarified. It should be noted that several of the genes whose expression is enhanced by PI3K signaling, i.e. blocked by Wortmannin, are known to be involved in resilience towards mitochondrial stress, like mitochondrially encoded ATP synthase membrane subunit 8 (mt-Atp8), and Mitochondria-encoded Cytochrome B (mt-Cytb), key enzymes of oxidative phosphorylation and generation of ATP (18,19), PI3K signaling also upregulates expression of cathepsin D (Ctsd), a regulator of apoptosis (20). On the other hand PI3K signaling does downregulate genes like Schlafen-2 (Slfn2) which has been shown to impact T cell quiescence and ultimately also protect from apoptosis (21,22), and also B cell lymphoma 2 (Bcl2), encoding a mitochondrial protein with established anti-apoptotic potential (23). PI3K signaling also impacts genes related to organization of the cytoskeleton and positioning of these cells, e.g. upregulating expression of cofilin (Clf1), a gene depolymerizing filamentous actin and of anti-apoptotic relevance (24), and integrin β2 (Itgb2), a component of LFA-1.

**Figure 3:**
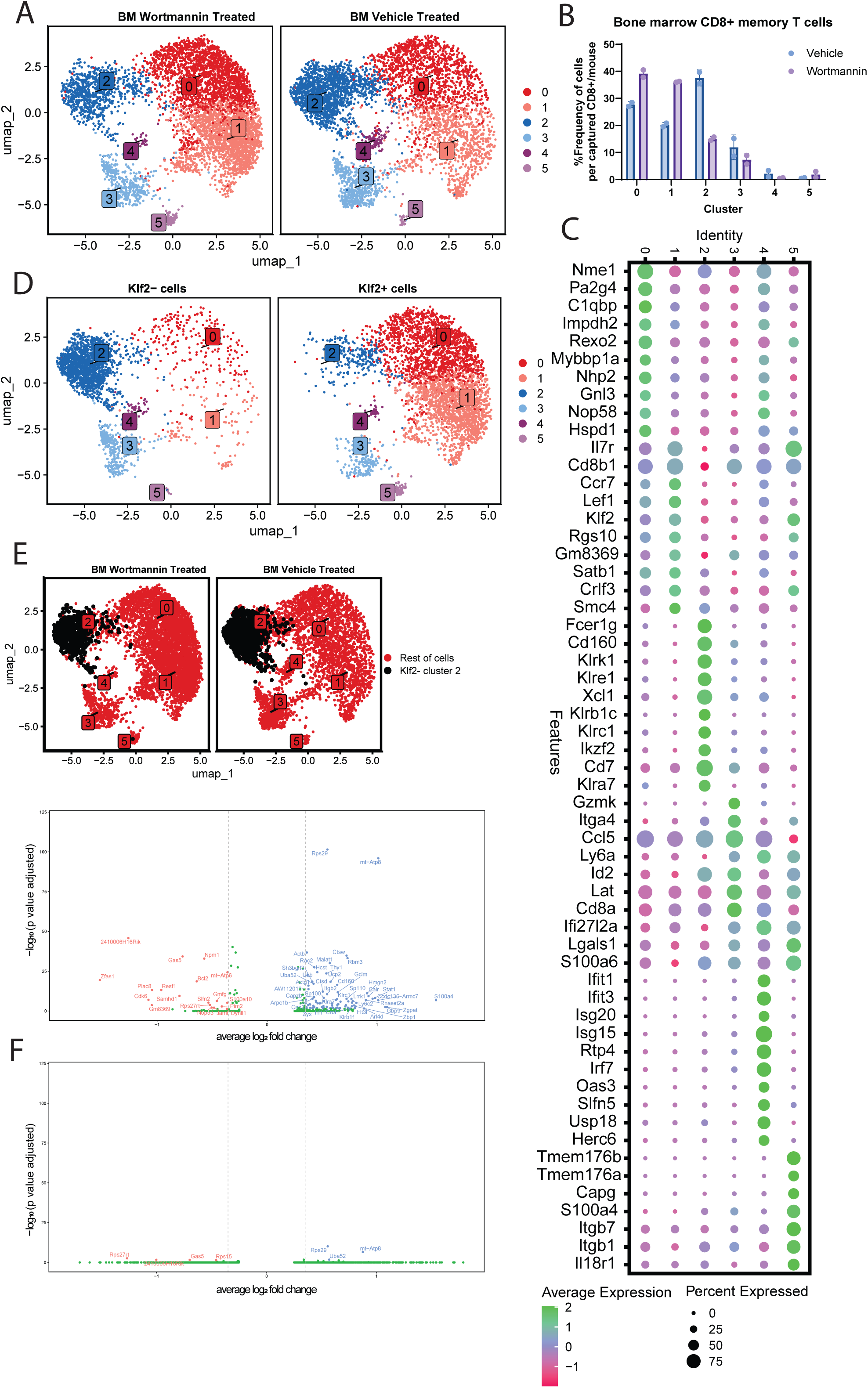
CD8 memory T cells isolated from bone marrow of mice treated either with 2.4 mg/kg body weight of Wortmannin or vehicle and submitted to single cell transcriptome analysis. Heterogeneity projected in UMAP with clustering based on second resolution from a Harmony integration (Ref) reveals 6 clusters (A). Frequencies of cells in each cluster are compared in (B), and signature genes shown in (C), based on top 10 most significant genes calculated in Seurat with default mean pct, log 2 fold change >=0.25 and adjusted p-value >= 0.05. Cells expressing Klf2 or not are shown in (D) independently of the treatment. In (E), Klf2-cells from cluster are highlighted in black for treated and vehicle-treated mice as they were used in the following volcano plot. In (F), differential transcription of Wortmannin-treated versus vehicle-treated cells is shown for surface CD69-positive Klf2-negative and for surface CD69-negative Klf2-positive cells of cluster 2, as indicated. For vehicle -treated mice, transcriptomes of 1006 Klf2-negative and 194 Klf2-positive cells, and for Wortmannin-treated mice, of 559 Klf2-negative and 55 Klf2-positive cells were compared. The differentially expressed genes and the frequencies of cells expressing them are shown in suppl. Figure 2.

In summary, these data show that KLF2 expressing memory T cells do not detectably use the PI3K signaling pathway for their persistence, when resting in the bone marrow, while KLF2-negative, surface CD69+ memory T cells of the bone marrow do so. Many of the genes regulated by PI3K are involved in the regulation of apoptosis, metabolism, mobility and positioning.

## Discussion

It is increasingly recognized, that memory T and B lymphocytes and long-lived memory plasma cells, the embodiment of immunological memory, are not only circulating in blood, but also residing in tissues (14,25–27). We previously demonstrated, that memory CD4 and CD8 T lymphocytes of the bone marrow are an essential component of long-term T cell memory for systemic antigens, both in mice and humans (14,15). In the murine bone marrow, they individually dock on to stromal cells and rest in terms of proliferation, transcription and mobility. This resembles the lifestyle of long-lived plasma cells, except that those secrete antibodies at very high rates (28). We and others have shown that long-lived plasma cells are maintained by integrin-mediated contact to the stromal cells, which is activating PI3K, inactivating Foxo1/3a, and preventing activation of caspases 3 and 7, i.e. metabolic apoptosis. In addition, survival of long-lived plasma cells depends on APRIL or BAFF signaling, activating the NF-kB pathway and preventing activation of caspase 12, i.e. anabolic apoptosis(6). Here we have tested the hypothesis, that tissue-resident memory T cells are persisting as well by integrin-mediated contact to the stromal cells they adhere to, and by integrin-induced PI3K signaling. We have blocked in mice *in vivo*, integrin-binding and PI3K, NF-kB or JAK/STAT signaling pathways, and evaluated the effect on CD4 and CD8 memory T cells, expressing CD69 on the cell surface or not, in spleen and bone marrow. Indeed, persistence of surface CD69-expressing memory T cells, both in bone marrow and spleen, was dependent on integrin-binding to stromal cells, and PI3K signaling. While persistence of surface CD69-negative memory T cells was dependent on integrins, they relied rather on NF-kB signaling than on PI3K signaling. Blocking Jak/Stat signaling did not affect memory T cells of the bone marrow, and only a fraction of memory T cells of the spleen.

In the experiments shown, we used the anti-integrin antibodies in the same concentration that previously had been shown to be effective for the depletion of bone marrow resident plasma cells(5). The other blocking agents were titrated out at ranges referring to their IC_50_ *in vitro*, in different cells. At the effective concentrations, those blocking reagents affected only subpopulations of memory T cells, allowing us to consider non-affected populations as specificity controls. Thus, the data presented here provide a consistent picture of the diversity of signaling pathways critically involved in the persistence of memory T cells in spleen and bone marrow.

Our data further question the relevance of homeostatic proliferation as the generic mechanism for the long-term persistence of T cell memory. The concept of homeostatic proliferation originally has been developed for memory T lymphocytes of the murine spleen, postulating that these cells are maintained by a homeostatic balance of cell death and proliferation, regulated by the cytokines, interleukin-7 and -15 (29–34). This is true in particular for bone marrow memory T cells and is based on the analysis of memory T cells from murine spleen, adoptively transferred, homing to several organs, including the bone marrow, and proliferating there with an apparent half-life of 14 days (35). The intuitive conclusion that homeostatic proliferation is the mechanism of persistence of T cell memory in general, was backed by experiments demonstrating the proliferation of endogenous CD8+ memory T cells in murine bone marrow, as determined by incorporation of Bromo-deoxy-Uridine (BrdU) (36,37) However, this result is in apparent contrast to the demonstration that almost all CD4+ and CD8+ memory T cells of murine and human bone marrow rest in G_0_ of cell cycle, i.e. are not proliferating, neither in terms of DNA synthesis, nor expressing Ki-67 (14,15,17), and that they also are not killed by cyclophosphamide, a drug targeting proliferating cells (38). We previously showed that BrdU-feeding induces the proliferation of CD8+ memory T lymphocytes it measures in the bone marrow, as indicated by induction of expression of Ki67, a protein not expressed by cells resting in G0 of cell cycle (39,40). Moreover, in humans, it has been shown that extended blockade of the JAK/STAT signaling pathway by Tofacitinib does not affect the numbers of circulating memory T cells (41). “Homeostatic proliferation” thus is confined to a subset of murine splenic memory T cells of unknown relevance.

Beyond “homeostatic proliferation”, persistence of memory T cells in tissues apparently relies on integrin-mediated binding to stromal cells, as we show here for the bone marrow and for CD4+ surface CD69+ memory T cells of the spleen. Interestingly, only in memory T cells expressing surface CD69, but not the transcription factor KLF2, i.e. classical “tissue-resident” memory T cells (11) persistence also critically depends on PI3K signaling. Persistence of surface CD69-negative memory T cells expressing KLF2 does not depend on the PI3K pathway. This pathway is not activated in the cells, because treatment with Wortmannin does not impact on their gene expression. Indicating that they use different pathways, one of which might be the NF-kB pathway. The impact of KLF2 on NF-kB signaling and on survival of T cells has been demonstrated before (42). As for the surface CD69+, KLF2-negative memory T cells, PI3K signaling for these cells apparently is necessary for survival and may be sufficient, whereas inhibition of NF-kB and JAK/STAT did not affect their persistence. We here describe the transcriptome signature of PI3K activation in a major cluster of CD8 memory T cells of the bone marrow, looking at the surviving cells. Of the 138 genes differentially expressed in the CD8 cluster 2 memory cells of Wortmannin-treated mice, several are of documented relevance in the context of apoptosis. It appears that PI3K has a multi-facetted impact on gene expression relevant for metabolism, apoptosis, positioning and mobility. Whether their differential regulation by PI3K reflects the transcriptome of the cells eventually ablated by Wortmannin, or whether it reflects gene expression of cells resilient to Wortmannin, remains to be shown, and is beyond the scope of the present analysis.

The diversity of molecular signaling pathways critical for persistence of memory T cells in spleen and bone marrow revealed here, may be resulting in different stabilities of persistence (43), and beyond bone marrow and spleen, these results provoke the question how memory T cells residing in other tissues, or circulating in blood and lymph are persisting.

**Limitations** of the present analysis are that the experimental approach of *in vivo* blocking will primarily discover those signaling pathways which are critical and non-redundant for the persistence of the cells in the bone marrow or spleen, like PI3K signaling, where our data show that this pathway is fundamental for persistence in the Klf2-positive memory T cells analyzed. Second, inhibitors not affecting all cells were not extensively titrated out *in vivo* further to determine whether they would affect all cells or just a subpopulation. This is particularly relevant for the Tofacitinib-sensitive cells, for which our previous analysis using cyclophosphamide had indicated that roughly 50% of CD8 memory T cells from the spleen were ablated, after 8 an 14 days, suggesting that only these 50% were dependent on cytokine-driven homeostatic proliferation (38). Thirdly, our data refer to persistence of the memory T cells in spleen and bone marrow. We cannot rule out formally that some of them left their organ of residency and survived outside of spleen and bone marrow. We consider this unlikely, since the critical involvement of the integrin-PI3K-Foxo1/3a pathway in the prevention of activation of caspase 3 mediated apoptosis of memory plasma cells has been demonstrated(6).

## Acknowledgements

The authors thank members of different labs from DRFZ for their valuable discussions. ESR was supported by the Mexican Ministry of Science, Humanities, Technology and Innovation (SECIHTI), through the CONACYT-DAAD 2021 stipendium for doctoral researchers. This work was supported by the BMBF through TReAT and CONAN and by the state of Berlin and the European Regional Development Fund through the grant EFRE1.8/11 and EFRE PersMedLab to MFM. ACS was supported by the Elsa Neumann stipendium. XD was supported by China Scholarship Council. This work was supported through the DGF Project-ID 375876048 and the Dr Rolf M. Schwiete Foundation to HDC.

## Materials and Methods

### Mice

All mice were purchased from Charles River, Germany, and maintained under SPF conditions at the mouse facility of the German Rheumatology Research Centre, Berlin. Experiments were performed according to institutional guidelines and German Federal laws on animal protection. Eight-week-old C57BL/6 mice were intraperitoneally immunized thrice at 21-day intervals with 100 μg 4-Hydroxy-3-nitrophenylacetyl coupled chicken gamma globulin (NP-CGG) emulsified in incomplete Freud’s adjuvant (IFA). Mice were then intraperitoneally administered with either Tofacitinib (20 mg/kg; Absource Diagnostics, München), Wortmannin (2,4 mg/kg; Absource Diagnostics, München), anti-LFA-1 (200 µg; Biozol Diagnostica, Hamburg) and anti-VLA-4 (200 µg; Biozol Diagnostica, Hamburg), rat IgG2a isotype control (200 µg; Biozol Diagnostica, Hamburg), rat IgG2b isotype control (200 µg; Biozol Diagnostica, Hamburg), IKK-16 (20 mg/kg; Absource Diagnostics, München), or Vehicle (DMSO) either on day 74, 76, and 78. After 2 days, mice were sacrificed on day 80. For single-cell sequencing experiments, mice were treated on day 74 and 76 with 2,4 mg/kg Wortmannin or vehicle and sacrificed on day 77.

### Flow cytometry

Single-cell suspensions were obtained from spleen and bone marrow. To generate a single-cell suspension of splenocytes, the spleen was homogenized by pushing it through a 70 µm cell strainer, filtered through a 40 µm strainer, washed, and centrifuged for 8 min at 300 x g and 4°C.

To generate a single-cell suspension from bone marrow mononuclear cells, the femur, tibia, and ilium were cut on one side and placed with the open end into a pierced 0.5 ml reaction tube that was put into a 1.5 ml reaction tube filled with 200 µl FACS buffer (PBS/0.1% BSA/2 mM EDTA). The reaction tubes were centrifuged for 15 seconds at 2900 x g and 4 °C. If bone marrow remained in the bone, the process was repeated. The bone marrow was pooled, filtered through a 70 µm cell strainer, washed, and centrifuged for 8 min at 300xg and 4 °C.

Before proceeding to the staining all cell suspensions were counted at the MACS Quant analyzer (Miltenyi, Bergisch-Gladbach). Cells were stained for 15 minutes at 4°C with anti-CD45 (REA737) anti-CD3 (eBio500A2), anti-CD4 (REA604), anti-CD8a/b (REA601/793), anti-CD44 (REA664), anti-B220 (REA755), and anti-CD69 (H1.2F3). Viability of cells was assessed by LIVE/DEAD Fixable Dead Cell Stain (Thermo Fisher Scientific, Waltham, MA). Stained samples were analyzed on a MACSQuant (Miltenyi, Bergisch-Gladbach, Germany). Flow cytometric data were analyzed by FlowJo software (FlowJo LLC, Ashland).

### Cell sorting

For sorting of CD4+ and CD8+ memory T cells, single-cell suspensions of bone marrow nuclear cells were prepared as described above in FACS Buffer containing 2 µg/µl Actinomycin D (Sigma-Aldrich Chemie GmbH, Taufkirchen), to freeze the transcriptional state. T cells were immunomagnetically sorted from the BM of mice treated with Wortmannin or Vehicle, according to the manufactureŕs instructions (Miltenyi). Enriched cells were incubated with Fc Blocking Reagent (Miltenyi) following manufacturer’s instructions and subsequently stained for 30 minutes at 4 °C with anti-CD3 (eBio500A2), anti-CD4 (REA604), anti-CD8a/b (REA601/793), anti-CD44 (REA664), and anti-B220 (REA755) in addition to the TotalSeq antibody cocktail. CD4+CD44+ and CD8+CD44+ memory T cells were sorted into tubes coated with PBS/BSA and mixed in a 1:1 ratio before continuing to the capture process for single-cell RNA sequencing.

### Single cell RNA-sequencing

For single cell library preparation, ex vivo FACS sorted CD3+CD4+CD44+DAPI- and CD3+CD8+CD44+DAPI-BM cells were applied to the 10X Genomics platform using the NEXT GEM 5’ v1.1 kit (10x Genomics) following the manufacturer’s instructions. Upon adapter ligation and index PCR, the quality of the obtained cDNA library was assessed by Qubit quantification, Bioanalyzer fragment analysis (HS DNA Kit, Agilent. The sequencing was performed on a NextSeq500 device (Illumina) using P3 Kit (100 cycles) with the recommended sequencing conditions (read1: 26nt, read2: 98nt, index1: 8 nt, index2: n.a.).

### Single cell transcriptome analysis

Illumina output was demultiplexed and mapped to the mm10 reference genome by cellranger-2.0.2 (10x Genomics Inc.) using refdata-cellranger-mm10-1.2.0 in default parameter setting and 3000 expected cells. Raw UMI-counts were further analyzed using R 3.5.2 and R 4.0.0 with Seurat (44,45) including log-normalization of UMI-counts, detection of variable genes and scaling. Uniform Manifold Approximation and Projection for Dimension Reduction and the underlying Principle Component Analysis was performed based on 50 components using variable genes and a perplexity of 50 as set by default.(46) Potential contamination cells lacking the expression of expressing *Cd3d* (CD45) and concomitant expression of *Cd79a*, *Vcam1*, *Csf1r* or *C1qa/b*, respectively were detected and excluded. Low quality cells were also excluded (47,48). The dataset was split into CD4+ and CD8+ Cells based on the CITESeq expression values of 1.8 and 1.3, respectively. After reanalyzing and reclustering the remaining datasets contained 6056 CD4+ cells and 10034 CD8+ cells. In order to differentiate between resident and non-resident cells the gene expression of *Klf2* and cell surface protein expression of CD69 was used.

---

**Supplementary Figure 1:**
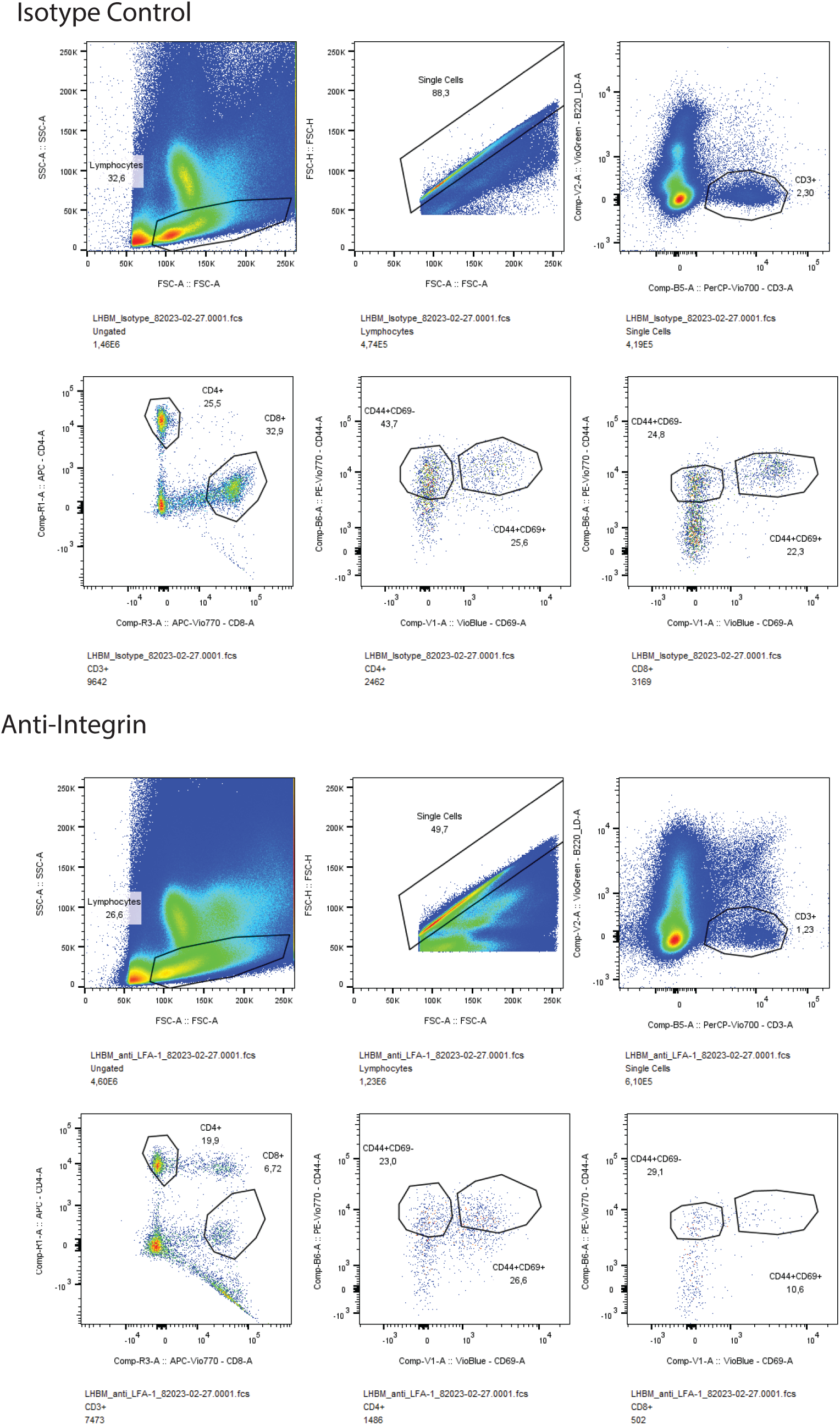
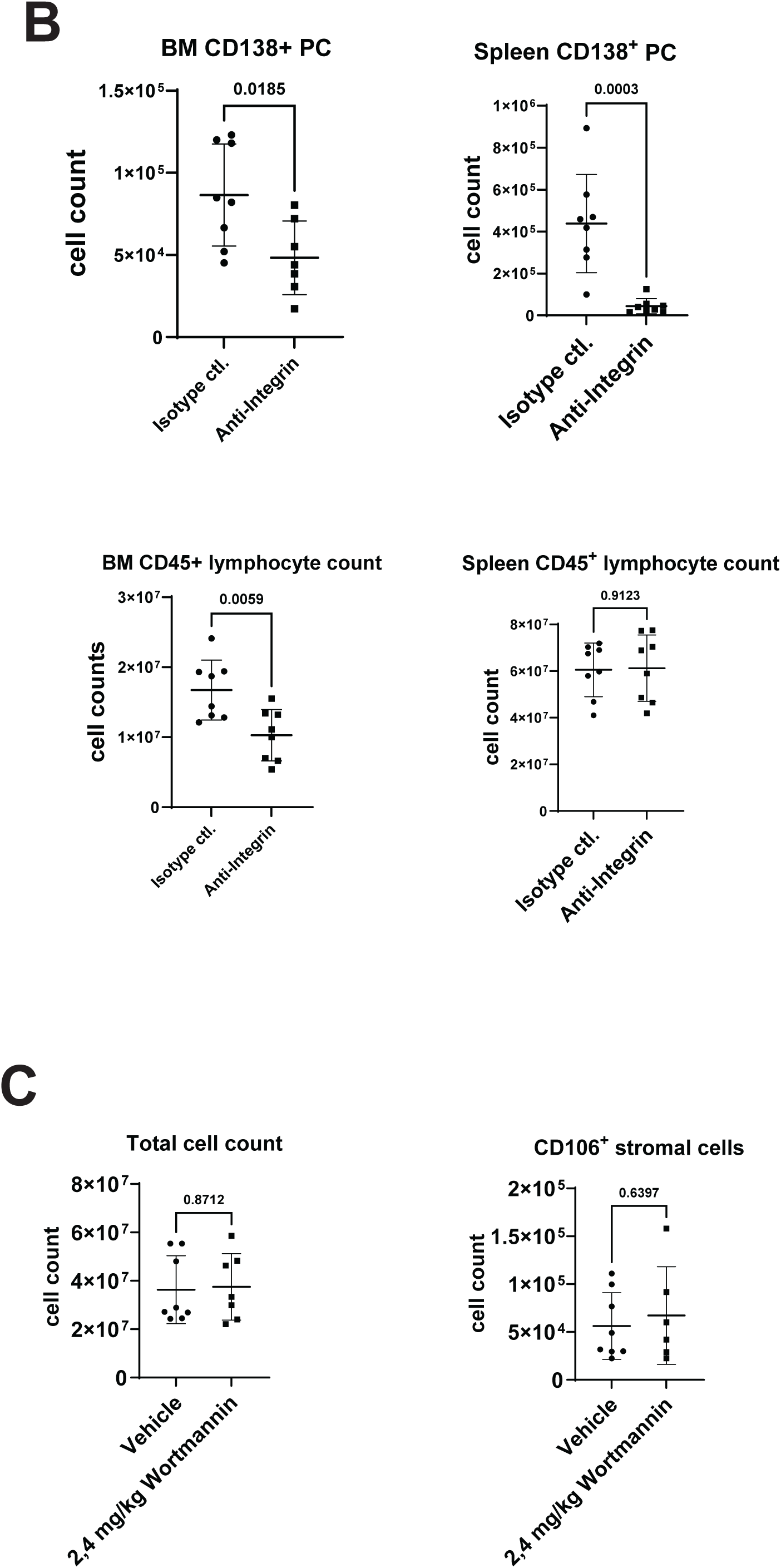
(A) shows the gating strategy exemplified for the bone marrow cells of mice treated with anti-integrin antibodies or isotype control antibodies. The depletion of plasma cells from bone marrow and spleen is documented in (B). Total cell counts of the bone marrow, and numbers of VCAM1+ stromal cells in (C). Mononuclear cells of bone marrow (BMMNC) were stained and analyzed at low flow rates. The absolute number of memory T cells was calculated as frequencies of the total bone marrow mononuclear cell count. Single viable T lymphocytes were gated in as negative for Zombie Aqua, and the B cell marker CD45R (B220), positive for CD3 and CD45, and either CD4 or CD8, with the scatter light profile of small lymphocytes. Memory T cells were identified according to expression of CD44 and CD69, as CD44^+^CD69^-^ and CD44^+^CD69^+^ memory T cells.

**Supplementary Figure 2:**
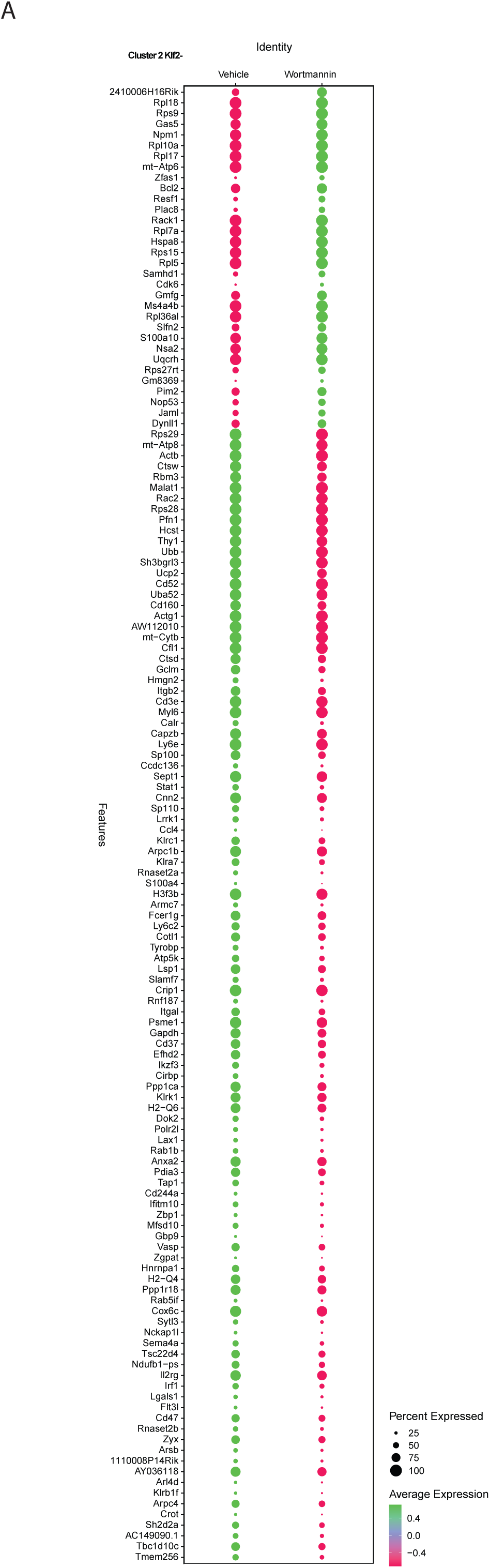
List of differentially expressed genes of surface CD69+, Klf2-negative CD8 memory T cells of cluster 2 of bone marrow. Color indicates up-(green) and downregulation (red), while the size of the circle indicates the frequency of cells of the cluster expressing the gene. The genes plotted were extracted from a differential gene expression analysis using Seurat and R based functions. Log 2-fold change threshold was >0.25 and adjusted p-value > 0.05.

**suppl. Figure 3.**
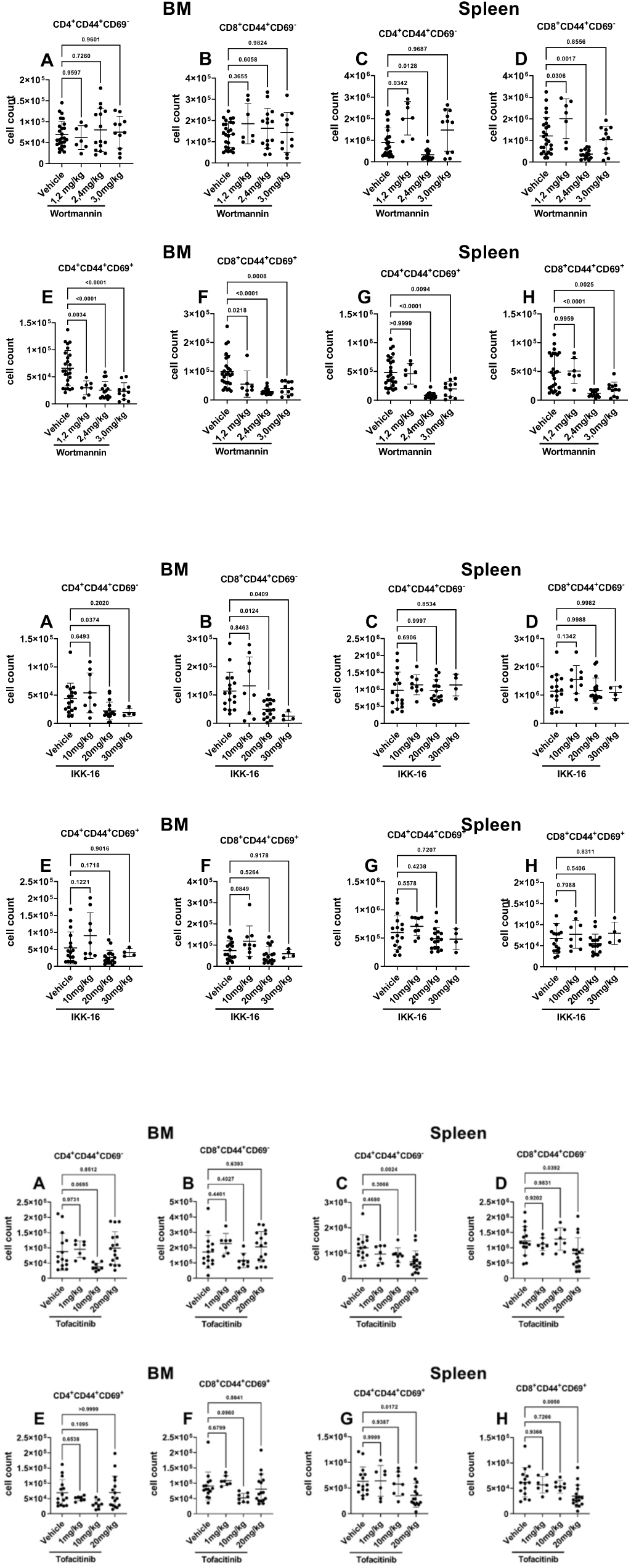

## Notes

### Competing Interest Statement

The authors have declared no competing interest.

